# The *Pgb1* locus controls glycogen aggregation in astrocytes of the aged hippocampus without impacting cognitive function

**DOI:** 10.1101/2023.11.22.567373

**Authors:** A. Gómez-Pascual, D. M. Glikman, H. X. Ng, J. E. Tomkins, L. Lu, Y. Xu, D. G. Ashbrook, C. Kaczorowski, G. Kempermann, J. Killmar, K. Mozhui, O. Ohlenschläger, R. Aebersold, D.K. Ingram, E. G. Williams, R. W. Williams, R. W. Overall, M. Jucker, D. E. M. de Bakker

## Abstract

In aged humans and mice, aggregates of hypobranched glycogen molecules called polyglucosan bodies (PGBs) accumulate in hippocampal astrocytes. PGBs are known to drive cognitive decline in neurological diseases but remain largely unstudied in the context of typical brain aging. Here, we show that PGBs arise in autophagy-dysregulated astrocytes of the aged C57BL/6J mouse hippocampus. To map the genetic cause of age-related PGB accumulation, we quantified PGB burden in 32 fully sequenced BXD-recombinant inbred mouse strains, which display a 400-fold variation in hippocampal PGB burden at 16-18 months of age. A major modifier locus was mapped to chromosome 1 at 72–75 Mb, which we defined as the *Pgb1* locus. To evaluate candidate genes and downstream mechanisms by which *Pgb1* controls the aggregation of glycogen, extensive hippocampal transcriptomic and proteomic datasets were produced for aged mice of the BXD family. We utilized these datasets to identify *Smarcal1* and *Usp37* as potential regulators of PGB accumulation. To assess the effect of PGB burden on age-related cognitive decline, we performed phenome-wide association scans, transcriptomic analyses as well as conditioned fear memory and Y-maze testing. Importantly, we did not find any evidence suggesting a negative impact of PGBs on cognition. Taken together, our study demonstrates that the *Pgb1* locus controls glycogen aggregation in astrocytes of the aged hippocampus without affecting age-related cognitive decline.

## Introduction

Polyglucosan bodies (PGBs) are formed by the aggregation of hypobranched glycogen molecules. Accumulation of PGBs is associated with neurodegenerative diseases including adult polyglucosan body disease (APBD) and Lafora disease (Duran et al., 2014). APBD is a rare neurogenetic disorder in which mutations in glycogen branching enzyme 1 (*GBE1*) reduce the efficiency of glycogen molecule synthesis, leading to hypobranched glycogen molecules prone to aggregation (Nitschke et al., 2022; Zebhauser et al., 2022). These aberrant glycogen chains aggregate in neurons and astrocytes, leading to cellular dysfunction across the peripheral and central nervous systems (Duran and Guinovart, 2015). As a result, APBD patients suffer severe peripheral neuropathy, muscle weakness and dementia. In Lafora disease, caused by mutations in epilepsy progressive myoclonus type 2 genes (*EPM2A* and *EPM2B*), PGBs accumulate in many tissues including the brain, skin, liver, heart, and skeletal muscle. Typically, this leads to ataxia and seizures in adolescence and the progressive development of severe dementia (Duran and Guinovart, 2015; Moreno-Estellés et al., 2023; Wang et al., 2007).

In addition to playing a role in neurological diseases, PGBs also arise in the aging brain under non-pathological conditions (Duran and Guinovart, 2015; Rai and Ganesh, 2019). These PGBs are most prominent in astrocytes of the hippocampus, but can also be found in other regions and cells of the periventricular and subpial regions of the aged human and murine brain (Duran and Guinovart, 2015; Jucker et al., 1994a, 1994b, 1992)

The astrocyte-specific accumulation of PGBs is likely due to the central role these glial cells play in the glycogen homeostasis of the brain (Palmer and Ousman, 2018). Astrocytes are a heterogeneous cell population with distinct functions, whose characterization is an ongoing process (Viana et al., 2023). For example, a subset of hippocampal astrocytes has recently been identified to show age-related autophagy dysregulation, leading to severely inflated autophagosomes. Although a direct relevance to cognitive function was not shown, these autophagy dysregulated astrocytes had detrimental effects on the number of synapses in adjacent neurons (Lee et al., 2022). Furthermore, astrocytic dysfunction could directly affect the age-related decline of hippocampal function, considering that hippocampal astrocytes play an active role in memory consolidation and the location-specific encoding of the reward system (Brazhe et al., 2023; Curreli et al., 2022; Denizot et al., 2019; Doron et al., 2022; Polykretis and Michmizos, 2022).

Here, we investigate the genetic architecture underlying misfolded glycogen aggregates in the aged hippocampus by analyzing the age-related occurrence of PGBs in 32 different BXD recombinant inbred mouse strains. One of the primary uses of the BXD family is to map genetic regions, also called quantitative trait loci (QTL), which influence phenotypic variation of a complex trait (Peirce et al., 2004; Taylor, 1978). Utilizing the BXD cohort and QTL mapping, together with novel behavioral and hippocampal transcriptomic and proteomic datasets, we identify a genetic locus controlling hippocampal PGB burden and evaluate the pathophysiological significance of these age-related glycogen aggregates.

## Results

### Polyglucosan bodies accumulate in autophagy-dysregulated astrocytes

To determine the cell type in which polyglucosan bodies (PGBs) accumulate, we validated the occurrence of PGBs in the hippocampus of 18-24 month old C57BL/6J males and female mice using antibodies targeting LBP110 (Jucker et al., 1994b). PGB burden was most pronounced in the CA1 and CA2 regions of the hippocampus but was also evident in all other hippocampal subregions including the dentate gyrus (**Figure 1A, B**). Double labeling with the astrocytic marker glial fibrillary acidic protein (GFAP) validated that PGBs are mainly localized to astrocytes and that a single astrocyte can contain a cluster of PGBs (**Figure 1C**), which is in good accordance with previous literature (Jucker et al., 1994b). Considering that the accumulation of glycogen might be linked to compromised autophagy (Aman et al., 2021; Kakhlon et al., 2021), we assessed whether PGBs arise in the recently defined autophagy-dysregulated astrocytes (APDAs) (Lee et al., 2022). Therefore, consecutive sections were stained with APDA marker SCRG1 and autophagosome marker SQSTM1, which both resulted in a staining pattern reminiscent of LBP110 (**Figure 1D,E**). Indeed, fluorescent co-labeling of LBP110 with SQSTM1 shows a strong overlap in the hippocampus (**Figure 1F-H**), indicating that hippocampal PGBs arise specifically in APDAs.

**Figure 1.**
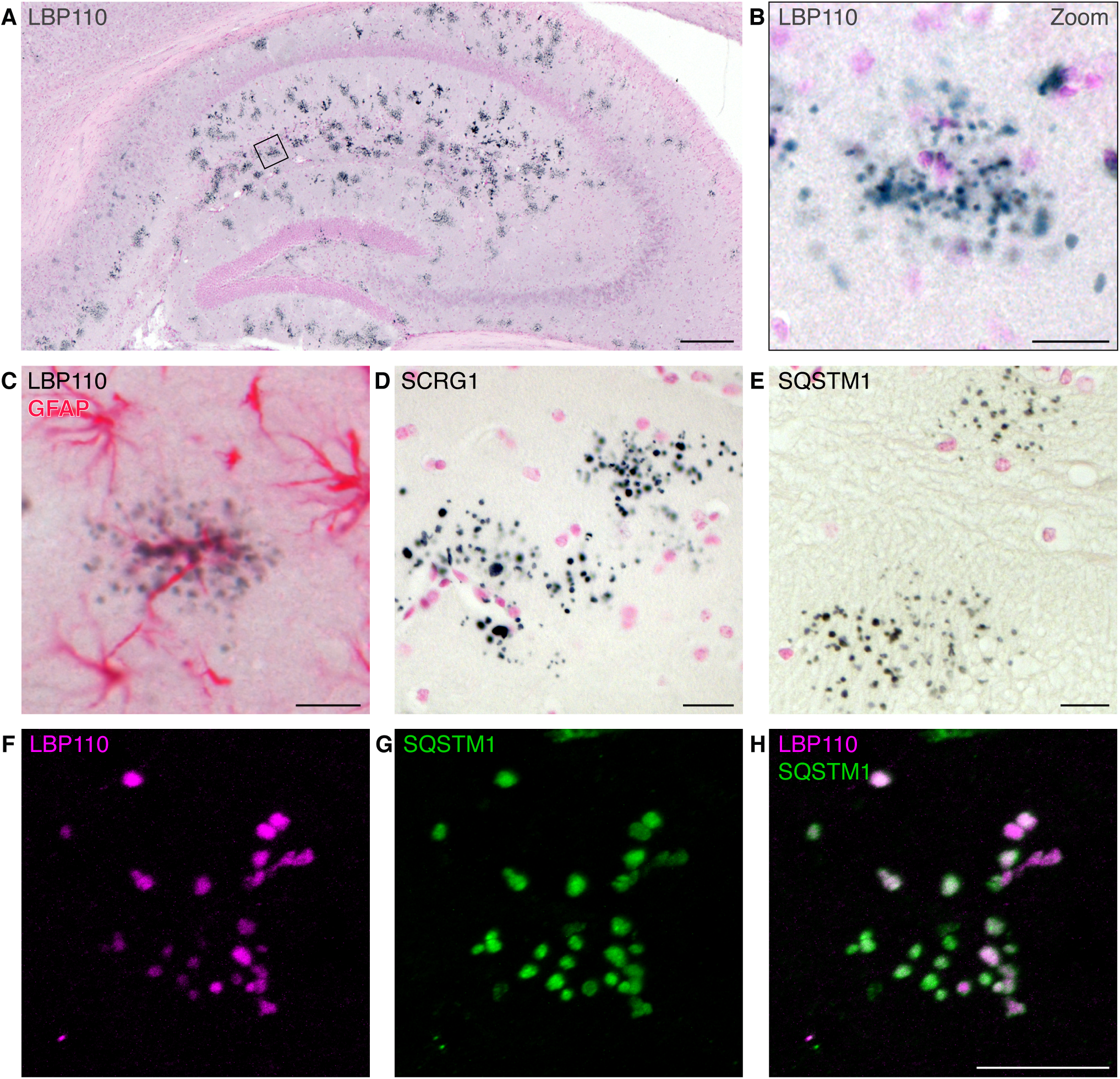
Polyglucosan bodies accumulate in the aging hippocampus. (**A,B**) The hippocampus of an 18-month old female, stained with an anti-LBP110 antibody targeting polyglucosan bodies. Scale bars represent 200 µm (A) or 20 µm (B). (**C**) Antibody staining to show astrocytes (GFAP, pink), and LBP110 (black) which marks polyglucosan bodies. (**D**) Antibody staining against SCRG1, a marker for autoph-agy dysregulated astrocytes. (**E**) Antibody staining against SQSTM1, an autophagosome marker. (**F-H**) Fluorescent antibody staining against SQSTM1 and LBP110, obtained from the dentate gyrus of a 2-year old male. Scale bars (C-H) represent 20 µm.

### Variation in hippocampal PGB burden maps to a gene-rich locus on chromosome 1

PGBs accumulate in the hippocampus of aged C57BL/6J (B6), but not DBA/2J (D2) mice (Jucker et al., 1994b). To investigate whether this heterogeneity extends to the BXD strains, we quantified the number of PGB clusters in the hippocampus of 142 aged females (16–18 months old) sampled from 32 BXD strains and both parental strains B6 and D2. The average number of PGB clusters, with a single cluster likely representing a single astrocyte, was determined from an average of four animals per strain (**Table S1**). Average PGB cluster counts varied over 400-fold among BXD strains, with a variation between 1.5 and 646 clusters per hippocampus (**Figure 2A**) (GeneNetwork dataset ID: BXD_10685). Extraordinarily high PGB burden was observed in B6 and BXD20 strains while PGBs were nearly undetectable in D2 and BXD32 strains (**Figure 2A-E**). Importantly, the variation of PGB burden between the inbred BXD strains allowed us to correlate PGB burden with several layers of information available for these strains, including their fully sequenced genomes (**Figure 2F**). We analyzed which genetic loci contribute to differences in PGB burden, which was mapped using sequence-based markers and a linear mixed model regression (GEMMA), which corrects for kinship (Ashbrook et al., 2021; Zhou and Stephens, 2012). The PGB cluster numbers among strains were strongly skewed and values were therefore log_10_ transformed prior to mapping (**Figure S1**) (GeneNetwork dataset ID: BXD_10686). The highest association was found on chromosome 1 (Chr 1) at 72.5 Mb (–logP 4.9, GCm38 assembly), with a 1.5 −logP confidence interval that ranged from 72.0 to 75.0 Mb (**Figure 3A-C**, **Figure S2**). Approximately, 25% of the variance in PGB burden can be explained by this locus (*r* = −0.51, *p* < 2.2 × 10^−3^, *n* = 34 strains), hereafter referred to as the *Pgb1* locus (**Figure 3D**).

**Figure 2.**
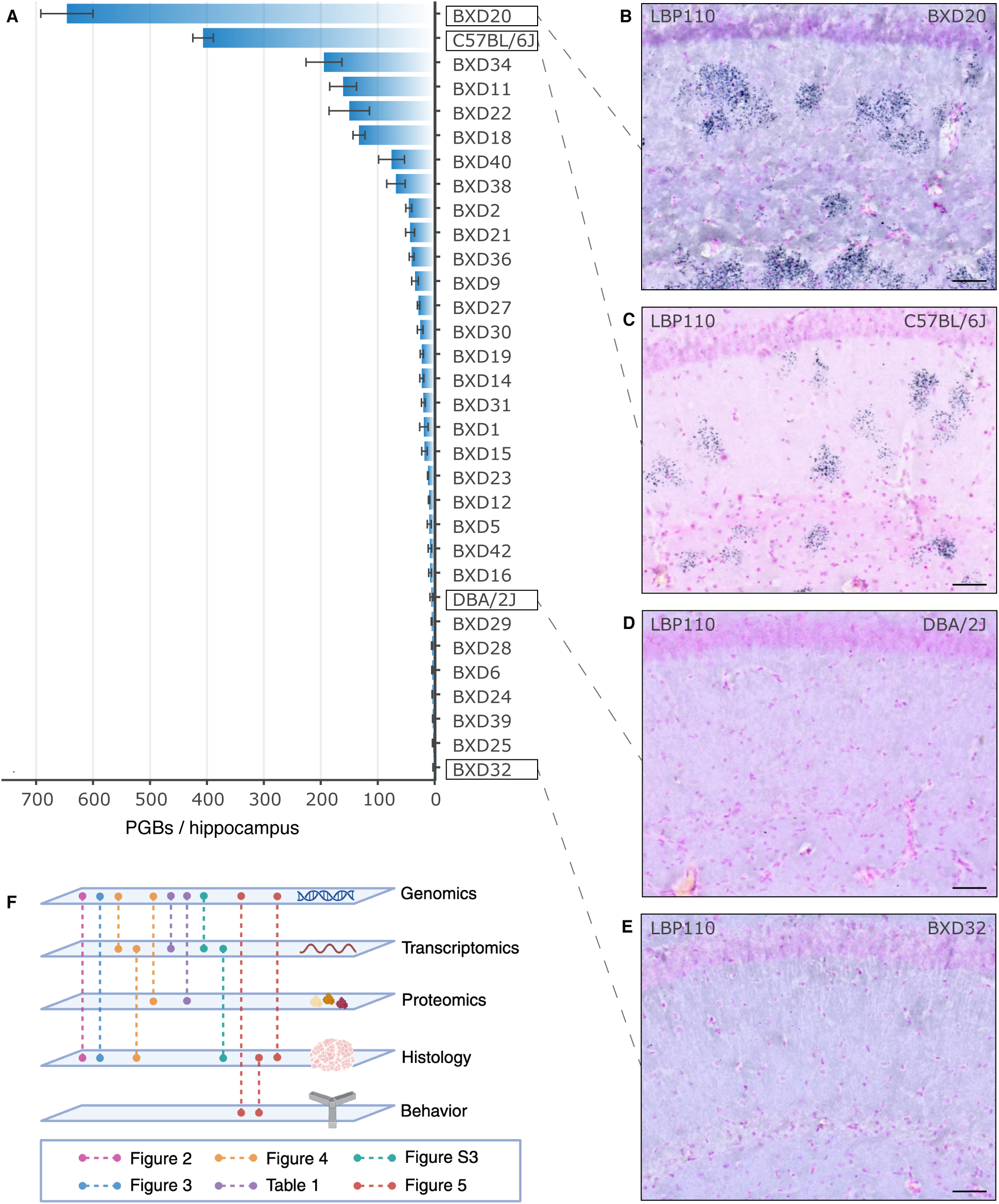
Polyglucosan body accumulation in the hippocampus varies widely between BXD stains. (**A**) Average number of PGBs per hippocampus of 32 BXD strains, as well as the parental strains C57BL/6J and DBA/2J. Genenetwork trait ID: BXD_10685. (**B-E**) Representative pictures of LBP110 staining in the hippocampus of the BXD32, C57BL/6J, DBA/2J and BXD20 strains. Scale bars represent 50μm. (**F**) Graphical abstract of the strategy used to determine the PGB phenotype.

**Figure 3.**
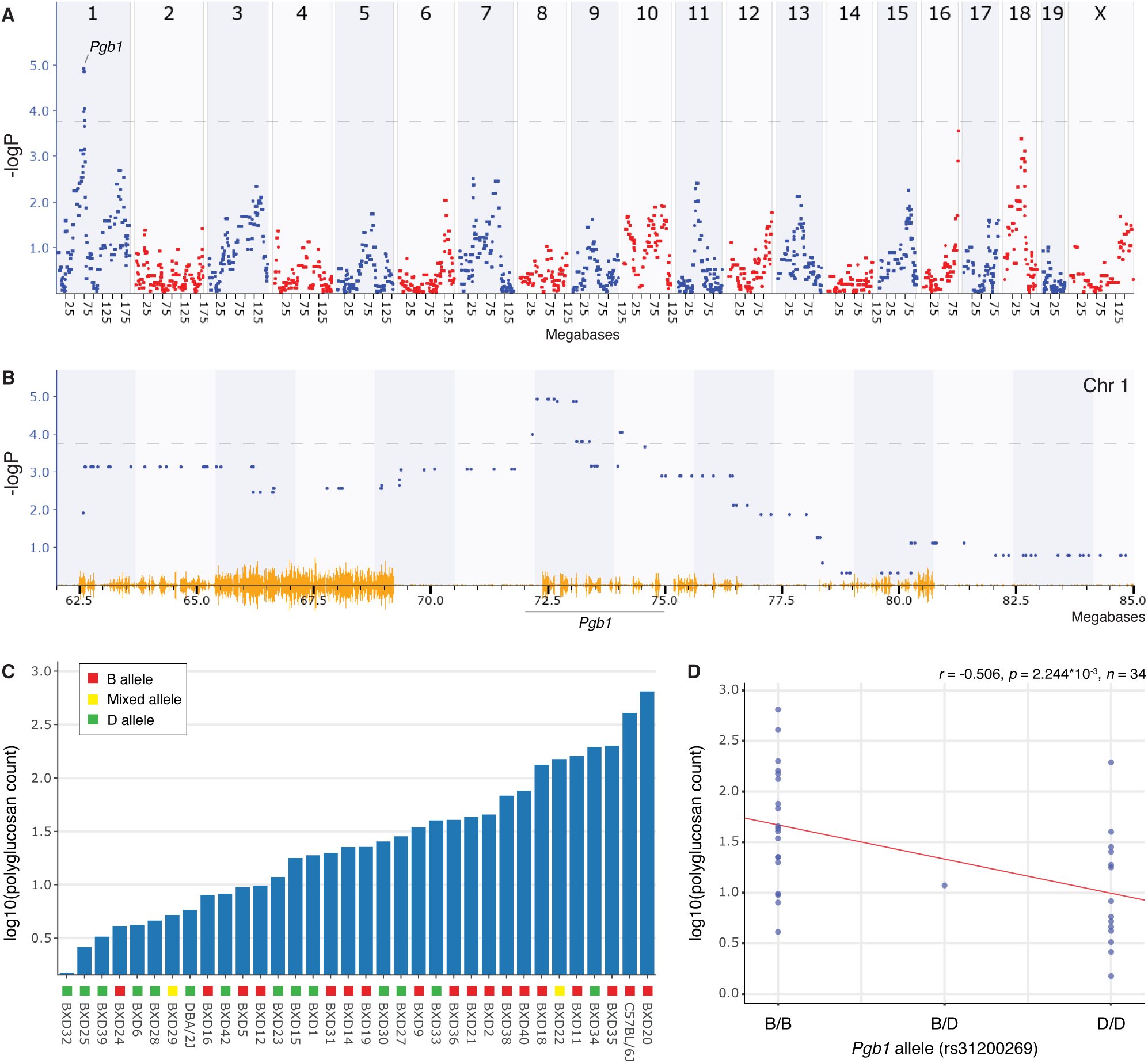
Quantitative trait locus mapping of polyglucosan body burden reveals a causal variation on Chromosome 1. (**A**) Manhattan plot indicating the correlation significance between polyglucosan body densities (GeneNetwork trait ID: BXD_10686) and linkage blocks across the genome. (**B**) Zoom-ins on the target region including the peak region of interest between 72 and 75 megabases (Mb). The yellow color indicates allele variations between BXD strains annotated at GeneNetwork.org. (A,B) The grey dotted line indicates genome wide significance at −logP of 3.832. (**C**) Log10 transformed average of PGBs per hippocampus of 32 BXD strains, as well as the parental strains C57BL/6J and DBA/2J. (**D**) Pearson collelation between the *Pgb1* allele (using SNP marker rs31200269) and PGB burden. The heterozygous strain (B/D) is BXD23.

### Candidate analysis identifies *Smarcal1* and *Usp37* as potential effectors of PGB burden

The *Pgb1* locus includes 94 potential coding regions of which 47 encode validated transcripts (42 protein coding, four microRNAs, and one long non-coding RNA; **Figure 4A**). To define allele variants most likely to modulate PGB density from within the *Pgb1* locus, we accumulated available data for each of the 42 protein coding genes (**Table S2**) (Ashbrook et al., 2022; Li et al., 2020; McLaren et al., 2016) and used specific selection criteria (**Figure 4A**).

**Figure 4.**
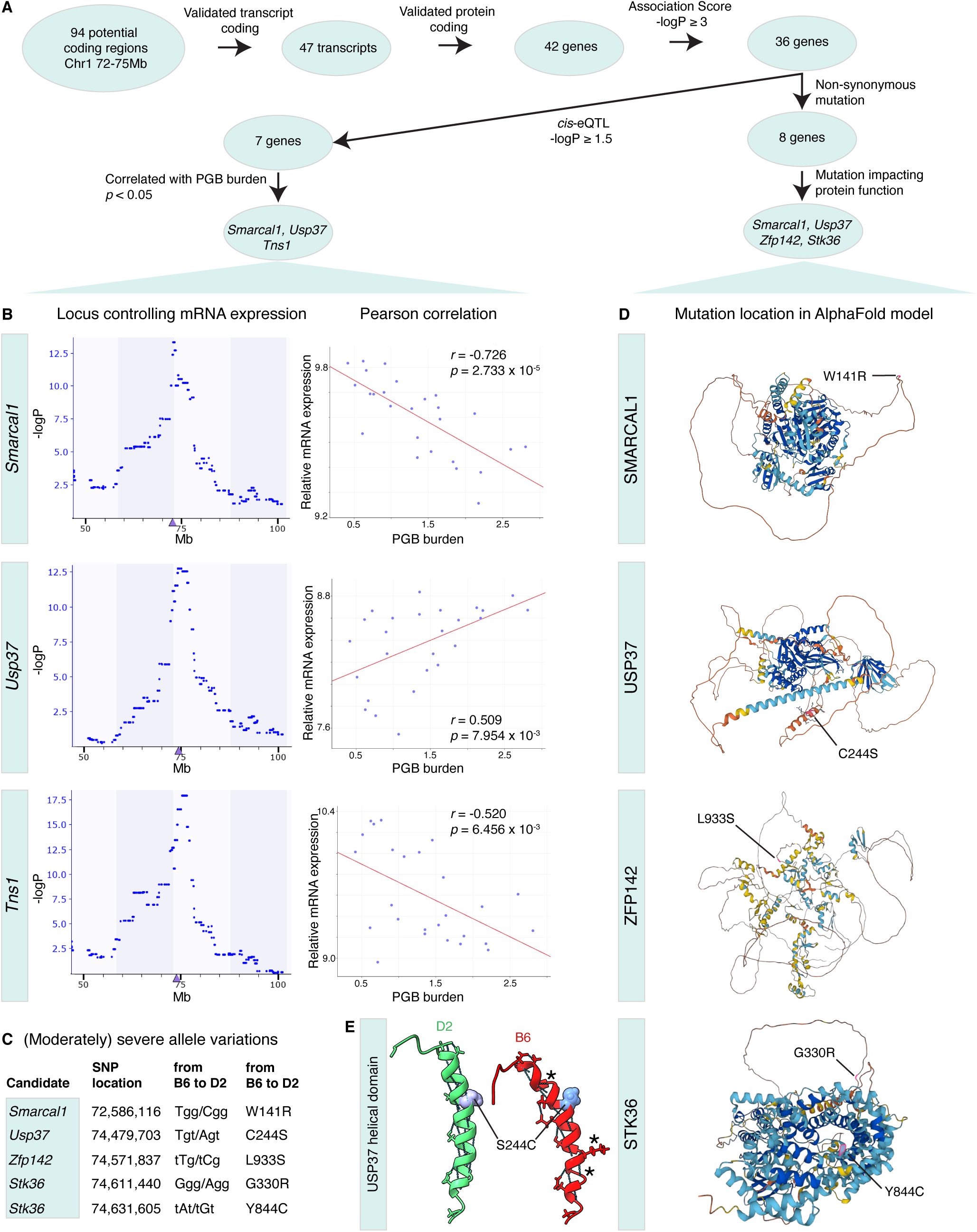
Selection of *Pgb1* candidate genes highlights potential impact of *Smarcal1*, *Usp37*, *Tns1*, *Zfp142* and *Stk36* on PGB burden. (**A**) Workflow of candidate selection, which was performed employing the data provided in Table S2. (**B**) *Cis*-eQTL peak location (left) and correlation of candidate gene mRNA expression with polyglucosan body burden (right). Pearson correlations were calculated from the mean values of 26 strains. (**C**) List of (moderately) severe allele variations between the *B* and *D* alelle within candidate protein coding genes (data extracted from Wang et al. 2016). (**D**) Alphafold models of the candidate genes, produced using the Ensembl browser and genome assembly GRCm39. Indicated are the locations of the allele variations (pink) between the B6 and D2 strains listed in (C). Colors correspond to AphaFold model confidence ranging from very high (dark blue) and confident (cyan) to low (yellow) and very low (orange). (**E**) Helical conformation in USP37 including the C244S allele variant which impacts the helical integrity at locations indicated by the asterisks. Blue lines indicate hydrogen bonds.

To begin, only protein coding genes with a linkage with PGB burden of −log*P* ≥ 3 (*n* = 36 protein coding genes) were considered. Variant gene candidates could influence PGB density via two distinct mechanisms, either by altering expression levels or by modifying protein function. To identify genes that exhibit altered expression levels due to allele variations within our target locus (a *cis*-regulatory effect), we produced extensive hippocampal exon array data from 235 BXD animals (223–972 days old; GeneNetwork dataset ID: GN392; **Table S3**). Analysis of mRNA expression-QTL (eQTL) was performed to identify the locus controlling expression level differences among BXD strains. Notably, we found that seven out of 36 candidate genes mapped as a *cis*-eQTL to the *Pgb1* locus (–logP ≥ 1.5) suggesting that sequence variations near the candidates (e.g., in promoters or enhancers) directly affects their expression level. We then selected those candidates that displayed a significant correlation (*p* < 0.05) between transcript expression levels and PGB burden and identified three candidates; *Smarcal1, Tns1,* and *Usp37* (**Figure 4B**). *Smarcal1* and *Tns1* were negatively correlated with PGB burden —correlations of −0.73 and −0.52, respectively— indicating a potential protective role for these genes against PGB burden. In contrast, *Usp37* exhibited a positive correlation with PGB burden (*r* = 0.509), suggesting a potential role in enhancing PGB numbers. Taken together, our analysis led to the identification of three candidate genes that may impact PGB burden through changes in gene expression levels.

To identify candidates that may affect PGB burden via changes in protein function, we focused on candidates with at least one non-synonymous mutation between the *B* and *D* alleles. From eight candidate genes that carried such mutations, we selected four candidates carrying variants with a previously reported high probability of affecting protein function (Wang et al., 2016): *Smarcal1*, *Usp37*, *Zfp142*, and *Stk36* (**Figure 4C**). We generated 3D protein structure models of these candidates to visualize the precise location of potentially high-impact gene variants (**Figure 4D**). Most allele variations could be found within flexible protein regions, while the USP37 allele variant (C244S) was located within a helical structure which might be compromised in the B6 variant (**Figure 4E**, asterisks). The localization of the identified allele variants in flexible and helical domains suggests that the binding affinity with interaction partners might be affected.

Interestingly, two candidates could affect PGB density through both changes in expression levels as well as through changes in protein function: the annealing helicase *Smarcal1* and the deubiquitinase *Usp37*.

### Phenome-wide association reveals *trans*-regulation of *Hp1bp3* mRNA expression by *Pgb1*

To evaluate whether the *Pgb1* locus has any *trans*-acting regulatory effects on the expression levels of mRNAs or proteins from genes located in different regions of the genome, we performed a phenome wide association study (PheWAS). SNP rs31200269 was used as a surrogate marker for the *Pgb1* locus since this SNP marker has the highest association with PGB density. SNP rs31200269 is located on Chr 1 at 72.626982 Mb (mm10 assembly) in intron 14 of *Smarcal1*.

We applied a PheWAS test with the aged hippocampal exon array data generated for 67 BXD strains and 234 mice which matched the age at which PGB burden was measured (mean age of 18 months). The strongest *trans*-acting regulatory effect of *Pgb1* was found with the expression of the terminal exon of heterochromatin protein 1 binding protein 3 (*Hp1bp3*) (probe set 10340721) (**Table 1A**). *Hp1bp3* is located on Chr 4, and is a modulator of cognitive function and conditioned fear memory in older BXD mice (Neuner et al., 2016). *Hp1bp3* expression in the aged hippocampus mapped to the *Pgb1* locus with a −logP of 6.3, showing higher expression in the *D* allele compared to the *B* allele (**Figure S3A-C**). PGB burden and *Hp1bp3* expression showed a negatively covarying trend (*r* = −0.34, *p =* 0.088, Pearson correlation) when correlating the data encompassing 26 BXD strains for which both mRNA and PGB cluster counts were available (**Figure S3D**). Expression of *Hp1bp3* increased as a function of age in a way consistent with linkage to PGB burden (**Figure S3E**; *r* = 0.29, *p* < 1 × 10^−5^) (Jucker et al., 1994b).

**Table 1A.** Trans-acting regulatory effects of *Pgb1* on mRNA expression in the hippocampus.

| Gene | Location (Mb) | Functional association<sup>a</sup> | −logP | B/D<sup>b</sup> | *r*<sup>c</sup> | Effect size | Record ID |
| --- | --- | --- | --- | --- | --- | --- | --- |
| *Hp1bp3* | Chr 4: 138.241495 | Heterochromatin organization | 6.31 | D | 0.38 | 0.094 | 10340721 |
| *Pacsin2* | Chr 15: 83.375607 | Intracellular vesicular transport | 3.29 | D | 0.28 | 0.050 | 10430997 |
| *Tspan3* | Chr 9: 56.131723 | Membrane complex stability | 3.39 | B | 0.35 | -0.039 | 10593842 |
| *Ext2* | Chr 2: 93.695631 | Glycosyltransferase active at the Golgi/ER | 4.19 | B | 0.20 | 0.072 | 10485225 |
| *Eid2* | Chr 7: 28.267881 | Negative transcriptional regulator | 3.19 | D | 0.33 | -0.030 | 10551483 |
| *Cd81* | Chr 7: 143.052795 | Membrane structure and regulation | 3.32 | B | 0.18 | 0.056 | 10559261 |
| *Fosb* | Chr 7: 19.302696 | Transcription factor complex | 3.16 | D | 0.57 | 0.080 | 10560481 |
| *Htr5a* | Chr 5: 27.841947 | Serotonin receptor | 2.55 | D | 0.47 | 0.048 | 10520355 |
| *Mtap2* | Chr 1: 66.175329 | Microtubule cytoskeleton organization | 2.68 | B | 0.45 | 0.032 | 10347036 |

<sup>a</sup>Functional information derived from UniProt and Gene Ontology (UniProt Consortium, 2023; Aleksander et al., 2023).
<sup>b</sup>Allele that increases expression. <sup>c</sup>Spearman correlation between PGB burden and mRNA expression.

To detect whether *Pgb1* controls PGB burden through changes at the protein level, we generated a hippocampal proteome dataset using 205 BXD females (6–24 months of age; sample details as in Williams et al., 2022) and performed a proteome wide PheWAS. We surveyed 17,799 proteins to identify those being trans-regulated by the *Pgb1* locus. A total of six proteins showed linkage scores at *Pgb1* with a −logP ≥ 2: GSTP2, UBA1, RHOB, ARHGDIA, PPP1R7 and NPEPPS (**Table 1B**). Taken together, nine transcripts and six proteins were identified to be trans-regulated by the *Pgb1* locus.

**Table 1B.**
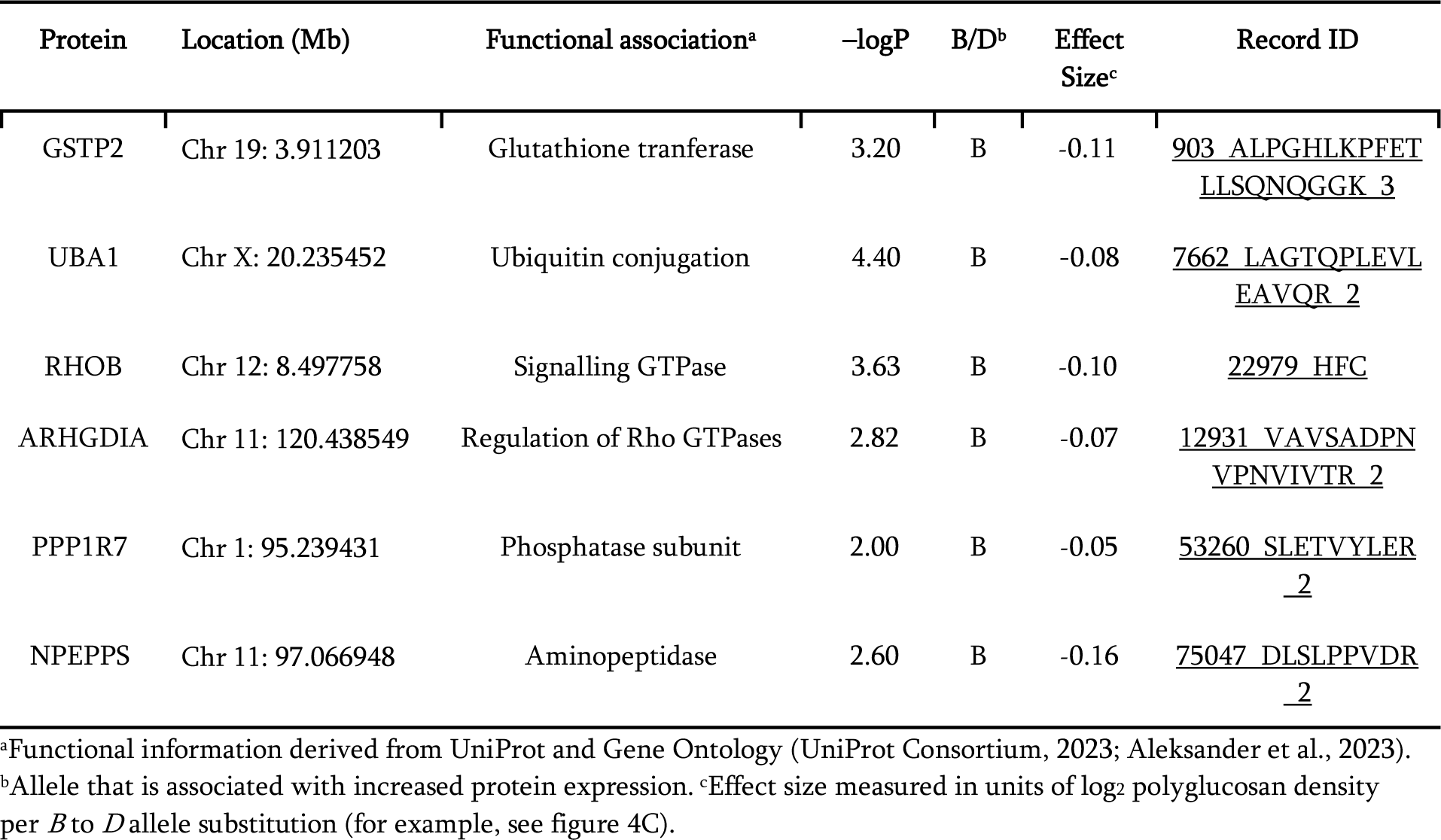
Trans-acting regulatory effects of *Pgb1* on protein expression in the hippocampus.

### PGB burden and age-relative cognitive decline appear uncorrelated

To assess the impact of PGB burden on cognition, the phenome-wide associations between *Pgb1* and all of approximately 12,000 phenotypes available at GeneNetwork.org (Mulligan et al., 2017; Sloan et al., 2016) were investigated. Notably, only two behavioral traits were linked to the *Pgb1* locus, including the level of activity in a Y-maze in 8-month-old females (*r* = −0.78, n = 23; GeneNetwork trait ID: BXD_20820; Neuner et al., 2019). This trait mapped to chromosome 1 with a −logP score of 4.8 at the *Pgb1* locus. Considering that the QTL peak association localized outside the *Pgb1* locus, the trait likely mapped due to a distal variant not associated with PGB burden directly. The *Pgb1* marker also covaried with cocaine-induced stereotypy (chewing, rearing; GeneNetwork trait ID: BXD_10315) in young adults (–logP = 4.2, *r* = 0.57, *n* = 23) (Jones et al., 1999), which is not directly linked with age-related cognition. Hence, we did not detect convincing evidence of an effector role for *Pgb1* on cognitive performance.

Transcriptomics were used to find potential correlations between the *Pgb1* locus and genes known to be involved in cognition. Differential expression analysis between *Pgb1 DD* and *BB* genotypes in mice older than 600 days showed that after adjusting the *p* values for multiple testing, only 12 differentially expressed genes (DEGs) were identified (*p* adj. < 0.05, **Table S4**). Seven of them are located within the *Pgb1* locus (**Figure 5A**). The small number of *trans*-regulated DEGs (n = 5) suggests that the downstream effects of the *Pgb1* locus on the aged hippocampal transcriptome is limited.

**Figure 5.**
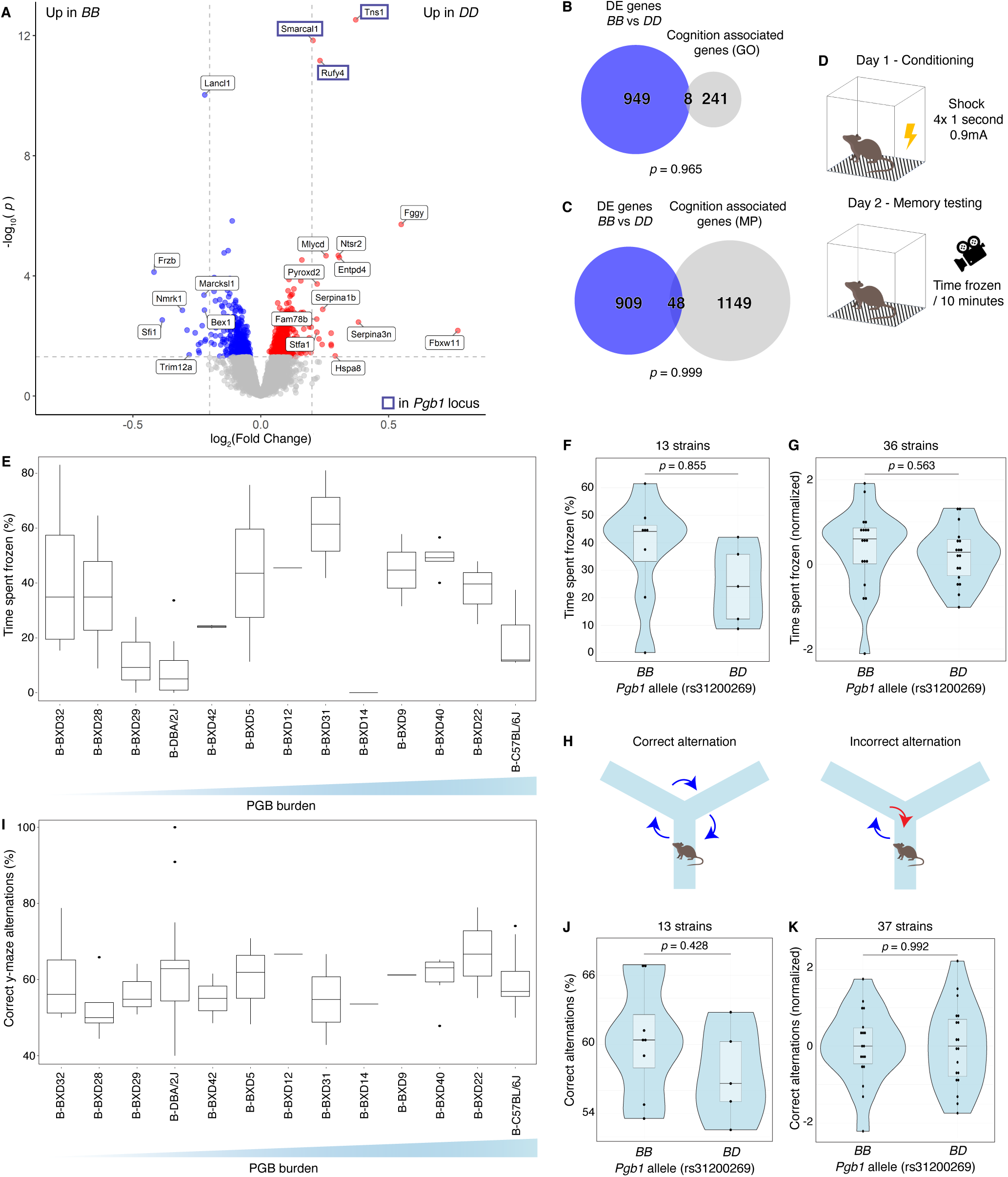
Cognitive capacity and PGB burden are uncorrelated in aged mice. (**A**) Vulcano plot indicating differentially expressed genes between old mice (age > 600 days) with a BB or DD allele at the *Pgb1* locus, determined using the R package limma (eBayes function). Note that the three highest overexpressed genes in DD are localized within the *Pgb1* locus (Highlighted by a dark blue box). (**B,C**) Differentially expressed genes (excluding genes located on the *Pgb1* locus) between old BB and DD carriers are not significantly enriched for genes known to be associated with cognitive performance, (B) GO:0050890_cognition and (C) MP:0014114_abnormal_cognition. (**D**) Workflow of conditioned fear memory test. (**E**) Time spent frozen plotted for each strain, ordered from low to high PGB density. All tested mice were 14-months of age and resulted from a backcross of homozygous BXD strains with the B parental strain. (**F**) Time spent frozen (%) by *Pgb1* allele. Means per strain (n = 13) were used to account for variable numbers of mice used per strain. No significant correlation was found between the Pgb1 allele and CFM score. (**G**) Pearson correlation plotted showing normalized conditioned fear memory score per allele at *Pgb1* (rs31200269). The data can be found under genenetworks trait ID 5XF_10474. In total, 142 animals of 14-months of age were tested, across 36 different strains. (**H**) Workflow of y-maze test. (**I**) Correct y-maze alternations plotted for each strain, ordered from low to high PGB density. The same mice were used as described in (G). (**J**) Correct y-maze alternations by *Pgb1* allele. Means per strain (n = 13) were used to account for variable numbers of mice used per strain. (**K**) Pearson correlation plotted showing normalized correct y-maze alterations per allele at *Pgb1* (rs31200269). The data can be found under genenetworks trait ID 5XF_10013. In total, 188 animals of 14-months of age were tested, across 37 different strains.

To exclude any effects that the *Pgb1* locus might have on the expression of genes correlated with cognition, we selected 995 DEGs using non-adjusted *p* values (< 0.05), of which only 35 showed a (−)logFC over 0.2. Among these DEGs, *Pgb1* locus genes *Tns1*, *Smarcal1* and *Rufy4* had the highest p-values (*P*<2.96×10^−13^, *P*<1.44×10^−12^, *P*<6.78×10^−12^, respectively) (**Figure 5A**). Next, we tested the overlap between the DEGs (excluding *Pgb1 cis*-regulated genes) and two cognition-associated gene lists from the Gene Ontology and Mammalian Phenotype repositories (GO:0050890_cognition and MP:0014114_abnormal_cognition, respectively) (**Figure 5B,C, Table S5**). Importantly, no significant overlap was found, which indicates that the *Pgb1* locus does not affect transcriptomic changes associated with cognition in mice over 600 days old.

To further investigate the potential correlation between PGBs and age-related cognitive decline, we performed conditioned fear memory (CFM) and Y-maze tests on 14-month-old BXDs from 42 different strains, which were backcrossed with C57BL/6J. At this age, PGBs are present in the B6 hippocampus and age-related behavioral alterations such as reduced CFM become apparent (Amelchenko et al., 2023; Hamieh et al., 2021; Jucker et al., 1994b; Wu et al., 2015). As the effect of the C57BL/6J genotype on PGBs requires two copies of the B6-allele, which was shown in a previously published B6/D2 intercross study (Krass et al., 2003), approximately half our cohort will carry a high risk (*BB*) genotype and half carry a low risk (*BD*) genotype.

No correlation was found between CFM (time spent frozen) and the PGB burden in the subset of overlapping strains (*r* = 0.0561*; p =* 0.855*; n* = 13 strains, GeneNetwork trait ID: 5XF_10474, **Figure 5D-F**). No significant correlation was found between *Pgb1* and CFM when considering all 142 animals across 36 strains (r = −0.100; *p* = 0.563; n = 36 strains), showing that animals with a *Pgb1 BB* allele, which are expected to carry more PGBs, do not underperform compared to mice with a *Pgb1 BD* allele (**Figure 5G**). We investigated the short-term spatial memory performance by Y-maze testing, for which we assessed the number of correct alternations defined as the subsequent investigation of previously uninvestigated arms of the maze (**Figure 5H**). No significant correlation was found between the percentage of correct alternations and PGB burden (*r* = 0.241*; p =* 0.428; *n =* 13 strains, **Figure 5I,J**). In addition, no significant correlation was found between *Pgb1* and Y-maze performance considering all 188 animals across 37 strains (*r* = −0.002; *p* = 0.992; *n* = 37 strains, GeneNetwork trait ID: 5XF_10013, **Figure 5K**).

Taken together the results of the phenome-wide association scans, the transcriptomic analysis and the CFM and Y-maze tests, we did not find any evidence suggesting a negative impact of PGBs on cognition.

## Discussion

Here, we show that glycogen aggregation and subsequent polyglucosan body burden in the hippocampus varies widely among BXD strains, which is in good accordance with previous studies that have shown that PGB abundance has a strong heritable component (Jucker et al., 1994a). One of the major advantages of using the fully sequenced inbred BXD strains is the availability of cross-correlation between independently produced datasets (Ashbrook et al., 2014; Chesler et al., 2004; Kempermann et al., 2006; Lu et al., 2001). Here, we generated two new datasets which can be utilized by the GeneNetwork community: mRNA and protein expression datasets for the aged BXD hippocampus. Using GeneNetwork.org and QTL-mapping, we mapped variation in PGB burden to a region on chromosome 1 (72–75 Mb) with a highly significant correlation (–logP = 4.9). The new transcriptomic dataset was used to identify two protein coding candidates likely to impact PGB burden, *Smarcal1* and *Usp37*. Both candidates harbor a non-synonymous mutation with the potential to impact protein function, and the mRNA expression levels of *Smarcal1* and *Usp37* significantly correlate with PGB burden. *Smarcal1* encodes an annealing helicase involved in repairing forked ends of DNA in response to DNA damage (Bansal et al., 2020) and *Usp37* encodes a deubiquitinase with limited functional characterization, although interestingly may also be linked to the DNA damage response (Wu et al., 2021), a key hallmark of aging (Schumacher et al., 2021). Considering that *Smarcal1* and *Usp37* have not been linked to glycogen metabolism in previous literature, follow-up studies should focus on testing the potential mechanistic link between these candidate genes and glycogen aggregate formation and degradation.

Recent work has identified the age-related occurrence of autophagy-dysregulated astrocytes (APDAs) in the hippocampus of C57BL/6 mice, which negatively affect synapse number and homeostasis of adjacent neurons (Lee et al., 2022). Here, we show that PGBs arise in cells displaying ADPA markers SCRG1 and SQSTM1. The finding that PGBs arise in APDAs suggests that the accumulation of PGBs in astrocytes might be a downstream effect of compromised autophagy, a primary hallmark of aging (Aman et al., 2021; López-Otín et al., 2023). While SCRG1/SQSTM1 antibody positivity defines ADPAs, whether or not these antibodies bind their intended epitope in hippocampal astrocytes remains to be determined, as PGBs are known to induce antibody reactivity in an epitope independent manner (Baglietto-Vargas et al., 2021; Jucker et al., 1992, Jucker et al., 1994b).

Hippocampal astrocytes are known to participate actively during memory consolidation and the location-specific encoding of the reward system (Brazhe et al., 2023; Curreli et al., 2022; Denizot et al., 2019; Doron et al., 2022; Polykretis and Michmizos, 2022). In addition, APDAs have been shown to reduce the number of synapses in adjacent neurons (Lee et al., 2022). Therefore, we hypothesized that the occurrence of PGBs in hippocampal astrocytes might have negative consequences on brain health and cognition during non-pathological aging. Importantly, we did not find any compelling evidence suggesting a negative impact of PGBs on cognition using phenome-wide association scans, transcriptomic analyses and CFM and Y-maze cognition testing of aged BXDs. The lack of association between PGB burden and cognition may be analogous to increased non-pathological aggregation of protein observed during aging, largely due to a decline in proteostasis (David, 2012). Emerging evidence suggests that insoluble protein aggregates, which are observed in aging as well as in many neurodegenerative disease states (Cuanalo-Contreras et al., 2023; Wilson et al., 2023), are not particularly neurotoxic. In contrast, small soluble misfolded oligomeric species are shown to contribute substantially to neurodegeneration (Rinauro et al., 2024). Our results, which indicate that the effects of glycogen aggregation on age-related cognitive decline are limited, support the notion that aggregation is not always detrimental to cognition and health.

In short, this study integrates multimodal data derived from the BXD mouse family to deepen our understanding of the accumulation of polyglucosan bodies in the aged hippocampus. PGBs were known to be detrimental to cognition in diseases such as APBD and Lafora disease, but whether this applied to age-related PGBs remained largely unknown. Here, we identify a novel locus controlling age-related PGB burden and highlight novel candidate genes and proteins implicated in glycogen aggregation. In addition, following extensive assessment of possible molecular and behavioral associations, we conclude that the physiological impact of PGB burden in these animals is likely unrelated to their cognitive performance.

## Materials and methods

### Animals and tissue preparation protocol for PGB assessment

Mice used for the assessment of PGBs were obtained as retired breeders between 1997 and 1999 from The Jackson Laboratory (Bar Harbor) and further aged at the Gerontology Research Center, NIA, NIH. A total of 144 female mice (all 17-19 months of age) were used: 32 BXD strains (n=133; 2 to 10 mice/strain; mean 4.1/strain; RRID:MGI:2164899), C57BL/6J (n=4; RRID:IMSR_JAX:000664) and DBA/2J (n=5; RRID:IMSR_JAX:000671). BXDs are progeny of crosses of female C57BL/6J (B6 or B) and male DBA/2J (D2 or D) parents. Animals were euthanized by pentobarbital overdose and perfused transcardially with 0.1 M phosphate-buffered saline (PBS) at pH 7.4. Brains were removed and immersion fixed in 4% paraformaldehyde, followed by 30% sucrose, and freezing in isopentane at −20C. Coronal sections throughout the hippocampus were cut on a freezing-sliding microtome at 25 μm. Visualization of the polyglucosan bodies (PGB) was done as previously described (Jucker et al., 1994b). In short, free-floating sections were rinsed with TBS (0.05 M Tris-buffer containing 1.5% NaC1, pH 7.4) and subsequently incubated in 0.3% Triton X-100 in TBS, followed by an incubation in 5% of goat serum in TBS. Sections were reacted for two days at 4°C with an antibody against a laminin-binding protein (LBP110) in TBS containing 2% blocking serum and 0.3% Triton X-100. Secondary antibody was goat anti-rabbit IgG. Antibodies were then detected by the ABC method with reagents from Vector Laboratories (Vectastain Elite ABC Kit, Vector Laboratories, Burlingame, CA, U.S.A.). The chromogen was diaminobenzidine, and the reaction product was intensified by adding NiCl.

### Quantification of PGBs

PGB are typically detected with periodic acid-Schiff staining (Jucker et al., 1994a, 1992; Manich et al., 2016). However, an antibody raised against the laminin-binding protein 110 (LBP110) was found to strongly bind PGB-lesions at light- and ultrastructural level and is largely used to identify PBGs in histological preparations (Jucker et al., 1994a, 1992; Kuo et al., 1996). The number of PGB clusters was assessed in a random systematic sampled set of every 10^th^ section through the entire hippocampus. Total number of clusters of LBP110-positive PGBs per unilateral hippocampus was estimated by counting all clusters PGB in all sections and multiplying the number with the section sample fraction (10 for the present analysis). The PGB numbers among strains were strongly skewed (GeneNetwork trait ID: BXD_10685) and values were therefore log_10_ transformed prior to mapping (GeneNetwork trait ID: BXD_10686).

### Hippocampal transcriptome in older BXD animals (GN712)

#### Animals

234 individual mice (140 females and 94 males) from 71 BXD strains, parental C57BL/6J, DBA/2J and D2B6F1 hybrid, were used in producing the hippocampal transcriptome dataset (GN712). These mice were obtained from UTHSC, ORNL, Beth Israel Deaconess or directly from the Jackson Laboratory. The age for a great majority of mice was between 12 and 28 months (average of 18 months). The animals were euthanized under saturated isoflurane, and brains were removed and placed in RNAlater prior to dissection. Cerebella and olfactory bulbs were removed; brains were hemisected, and both hippocampi were dissected whole in Dr. Lu’s lab. Hippocampal samples are close to complete (see Lu et al., 2001) yet may include variable amounts of subiculum and fimbria. All procedures were approved by the UTHSC Institutional Animal Care and Use Committee.

#### RNA Extraction and data generation

RNA was extracted using the RNeasy mini kit (Qiagen, Valencia, CA, USA) according to the manufactures’ procedure. RNA purity and concentration was checked using 260/280 nm absorbance ratio, and RNA integrity was analyzed using the Agilent Bioanalyzer 2100 (Agilent Technologies). Samples with RNA Integrity Numbers (RIN values) > 7.5 were run on Affy MoGene1.0 ST at the UTHSC. This array contains 34,700 probe sets that target ~29,000 well-defined transcripts (RefSeq mRNA isoforms). A single probe set is a collection of about 27 probes that target known exons within a single gene. The multiple probes design provides a more comprehensive coverage of transcripts from a single gene.

#### Data processing

Probe set level intensity values were extracted from the CEL files using the Affymetrix GeneChip Operating Software (RRID:SCR_003408). Data normalization was performed as previously described (Geisert et al., 2009), using the R package “affy” (RRID:SCR_012835) (Gautier et al., 2004). The Robust Multichip Averaging protocol was used to process the expression values. The array data were then log transformed and rescaled using a z-scoring procedure to set the mean of each sample at eight expression units with a SD of two units.

### Quantitative trait mapping

Two datasets were generated. The original untransformed dataset for 32 members of the BXD family (GeneNetwork trait BXD_10685) had a range of values from almost free of any polyglucosan aggregates (1.5 PGBs per hippocampus in BXD32) to very high densities of PGB (646 PGBs per hippocampus in BXD20). Due to the extreme skew (3.3) and kurtosis (12) the original data were not well suited for mapping (Figure S1). In contrast, log_10_ transformed data (GeneNetwork trait BXD_10686) were suitable for mapping and have low skew (0.29) and kurtosis (–0.58) and mean and median are both about 1.4. (Figure 1D). We used existing whole-genome sequence (WGS) marker maps (Ashbrook et al., 2021; Sasani et al., 2022) and the GRCm38 (mm10) mouse assembly. These high density genetic maps were used in combination with GEMMA, a linear mixed model method which corrects for differences in relatedness among strains (Zhou and Stephens, 2012). We mapped using a set of 20 kinship matrixes, each computed by excluding all markers on one of the 19 autosomes or the X chromosome (Yang et al., 2014). Support intervals are given by the −2.0 logP drop from the high point. Effect sizes are uncorrected for the Beavis effect but do not include parental phenotypes (Beavis, 1994; Beavis et al., 1991; Xu, 2003).

### Selection of data on candidate genes shown in Table S2

In Table S2, we list key data on coding genes within the *Pgb1* locus.

#### Mapping mRNA expression

We mapped every gene on the locus by itself from the “UTHSC BXD Aged Hippocampus Affy Mouse Gene 1.0 ST(Sep 12) RMA Exon Level’’ dataset provided in genenetwork.org (GN Accession: <u>GN392</u>). In this process, we chose between multiple results per gene based on the number of represented strains overlapping with the PGB strains, standard error being low, effect sizes being high and highest possible association scores. We noted the mean mRNA expression, given as a normalized Z-like scores value with a mean value of eight units with a SD of two (Geisert et al., 2009) in hippocampal array data, to confirm the gene being represented in our tissue of interest. Values lower than seven were considered to represent negligible expression. *cis-*eQTLs using microarray data can contain false positives due to higher binding affinity of the B allele, for which the microarray probes were designed. We reduced the possibility of the *cis-*eQTL being a probe binding artifact, by verifying its existence in other tissues as well, following the assumption of the *cis-*eQTL effect being predominantly conserved over tissues (Westra and Franke, 2014). Furthermore, we have prioritized candidates without proven irregularities in array sets found in mismatched SNP alleles on the UCSC Genome Browser (RRID:SCR_005780; https://genome.ucsc.edu/) (Kent et al., 2002) for mouse genome (GRCm38/mm10), another redirection given by genenetwork.org to verify probes via the “BLAT” tool (Kent, 2002).

#### Identification of allele variations affecting protein function

Proteomics data on the BXD family provided by David Ashbrook (Ashbrook et al., 2022) were used to investigate predicted protein altering segregating variants and whether those would be predicted to be deleterious variants according to the Ensembl Variant Effect Predictor (RRID:SCR_007931) (McLaren et al., 2016), providing another category measuring the severity of consequences of a gene variation in the family. Subsequently, we examined genes with a high number of mutations based on (Wang et al., 2016), which indicates the mutation severity based on the maximum Grantham score.

#### Age associated gene expression changes

Age-associated gene expression changes were quantified using bulk RNA-seq data from C57BL/6J male mouse hippocampi across the lifespan (2 months, 12 months, 24 months, n=3 per timepoint) (Li et al., 2020). Raw paired end sequencing reads were quality assessed and processed through fastp (v0.20.1; RRID:SCR_016962; Phred quality > 40; 10% unqualified bases permitted) (Chen et al., 2018). Processed reads were mapped and quantified using Salmon (v1.10.1; RRID:SCR_017036) in mapping-based mode with seqBias, gcBias and posBias parameters enabled (Patro et al., 2017). A decoy-aware transcriptome assembly (concatenated genome and transcriptome) used for mapping was derived using *Mus musculus* GRCm38 reference files (release 102 from Ensembl). Gene level expression estimates were achieved using tximport (v1.22.0; RRID:SCR_016752) (Soneson et al., 2015) and normalized counts calculated using DESeq2 (v.1.34.0; RRID:SCR_015687) (Love et al., 2014).

#### Assessing cell type specific expression

Single-cell RNA-seq data was retrieved from the Allen Brain Atlas portal (RRID:SCR_002978), to assess cell-type specific expression of the candidate genes (Yao et al., 2021).

### Structure prediction

Structure predictions of Smarcal1, Usp37, Zfp142, and Stk36 were performed using either the AlphaFold model on the web browser of Ensembl (release 112, Harrison et al., 2024) (Figure 3D) a local AlphaFold (Jumper et al., 2021) implementation (employing python v.3.10.13; hmmer suite v.3.4; hhsuite v.3.3.0; kalign v.3.4.0; ptxas v.11.7.64)(Figure 3E). Calculations were executed on computer servers with AMD EPYC 7F52 16-core processors. Modeling was run using default options (e.g. full genetic database configuration) with five seeds for each of the five models. The AlphaFold confidence score was employed to rank the predictions.

### Differential expression analysis in older BXD animals

To detect a possible relationship between PGB burden and cognition in aged hippocampus, a differential expression analysis was carried out between Pgb1 DD and BB alleles in mice older than 600 days. To this end, transcriptome data from a total of 72 mice, 36 with DD and 36 with BB alleles were used (GN712). Only protein coding genes were kept for downstream analysis (n=15876). First, a linear model for every transcript of the expression matrix was created with the lmFit function from the limma R package (Ritchie et al., 2015), which performs a t-test in the case of a two-class comparison. Then, statistics of differential expression analysis (moderated t-statistics, moderated F-statistic, and log-odds) were computed by the empirical Bayes method through the eBayes function from the same R package. DEGs were represented in a volcano plot. To investigate whether the DEG list (with non-adjusted p values) was enriched for cognition associated genes, the intersections between the DEGs and two independent cognition related gene lists (GO:0050890_cognition and MP:0014114_abnormal_cognition; Aleksander et al., 2023; Smith and Eppig, 2009; Table S3) were tested through a Fisher’s exact test using testGeneSet from the CoExpNets R package (Botía et al., 2017).

### PGB burden impact in cognitive performance

To account for the potential influence of accumulation of PGBs in hippocampal astrocytes on learning and memory functions, a search was proposed using the dataset: “UTHSC BXD Aged Hippocampus Affy Mouse Gene 1.0 ST (Sep12) RMA Exon Level” (GN Accession: GN392). The search was “rif=learning”, where rif = reference into function—an NCBI summary of gene functions with links to references was used. The list of results was sorted by peak location and focused on the genes that map to 72–75 Mb. Finally, mapping was repeated using GEMMA to validate the results.

### Phenome-wide association analysis of *Pgb1* locus

Two SNP markers within *Pgb1* were selected: (1) a proximal marker, rs31200269, at 72.62 Mb and (2) a distal marker, rs13475923, at 74.07 Mb. These markers are in strong linkage (*r^2^* = 0.79) and only two strains among those phenotyped differ in their genotypes—BXD23 and BXD29. Both proximal and distal markers were used as proxies for the *Pgb1* locus and in the phenome-wide association analysis to find traits that a) covary with these markers and b) that map within the *Pgb1* locus with linkage scores (–log*P* values) higher than 2.0. Candidate phenotypes, including expression traits, were considered of high interest when they were associated with behavioral differences or with the metabolism of polyglucosan or the formation of other types of aggregates in CNS.

### Contextual Fear Conditioning

Studies have shown that CFM is an effective readout of hippocampal function in spatial memory and learning confidence (Amelchenko et al., 2023; Hernández-Mercado and Zepeda, 2022) and have been demonstrated to deliver readable outcomes in cognitive performance tests in an age-related context (Hamieh et al., 2021; Hernández-Mercado and Zepeda, 2022; Verbitsky et al., 2004; Wu et al., 2015). Contextual fear conditioning (GN trait ID: 5XF_10474) was performed as previously described (Neuner et al., 2019). This was used to characterize cognitive function across the B-BXDs at either 6 or 14 months of age. We used F1 crosses between the C57BL/6J and BXD strains (B6-BXDF1s), as these were being phenotyped in a parallel project. On the first day of training, mice were placed in a training chamber and four 0.9 mA 1 s foot shocks were delivered after a baseline period of 180 s and then repeatedly after an interchangeable interval of 115 ± 20 s. Four post-shock intervals were defined as the 40 s following the offset of each foot (Colbourn Instruments, PA, United States). The percentage of time spent freezing following the final shock was used as a measure of contextual fear acquisition across the panel. Twenty-four hours after training, mice were placed back into the training chamber and the percentage of time spent freezing throughout the entire 10-min test was measured as an index of contextual fear memory. A DeepLabCut model was trained to recognize 13 points on a mouse, following the labelling system of Sturman et al., 2020 DLCAnalyzer was used to measure freezing events (Sturman et al., 2020).

### Y-Maze

A Y-maze test (GN trait ID: 5XF_10013) in F1 crosses of BXDs and non-transgenic C57BL/6J was used to assess short-term spatial memory performances in a novel environment. Sessions were conducted over 10 minutes, during which the performance was recorded using a Raspberry Pi camera operating at 30 frames per minute (fps). Video frames were labeled and trained using DeepLabCut (RRID:SCR_021391) (Mathis et al., 2018; Sturman et al., 2020) to track animals in the maze. Alternation behavior was assessed using a R script integrated with DLCAnalyzer. A mouse was counted to have entered an arm of the maze when the tail base crossed into that region. Mice exhibiting fewer than seven total alternations throughout the testing were excluded from further analysis. A correct alternation was recorded when a mouse exited one arm and subsequently entered a new arm, not last explored. A percentage of incorrect and correct alternations has been computed out of the recorded data on a single subject basis.

## Data Availability

All data used are publicly available. The overview of the used datasets and where to find them are in **Table S3**.

## Acknowledgements

We would like to thank the Summer School *Systems Genetics of Neural Ageing* for bringing us together and spurring our international collaboration. We would also like to acknowledge the funding for the Summer School 2022 from the e:Med Systems Medicine programme of the BMBF (*Bundesministerium für Bildung und Forschung*; German Ministry of Education and Research) to RWO. In addition, we would like to thank the FLI Imaging core facility for their assistance. Alicia Gómez-Pascual is supported by Fundación Séneca, Región de Murcia, Spain (21259/FPI/19). D.E.M. de Bakker is financed by a Rubicon scholarship (452021116) from the Dutch Research Council (NWO). NIH NIA R01AG070913-01 (Williams/Johnson), R01AG075813-01 (Ashbrook) and R01AG075818 (Kaczorowski). We acknowledge the help of Larry Mobraaten (Jackson Laboratory, Bar Harbor, MN) with the BXD strains. This research was funded in part by Aligning Science Across Parkinson’s [ASAP-000509] through the Michael J. Fox Foundation for Parkinson’s Research (MJFF). For the purpose of open access, the authors have applied a CC BY public copyright license to all Author Accepted Manuscripts arising from this submission.

---

**Figure S1.**
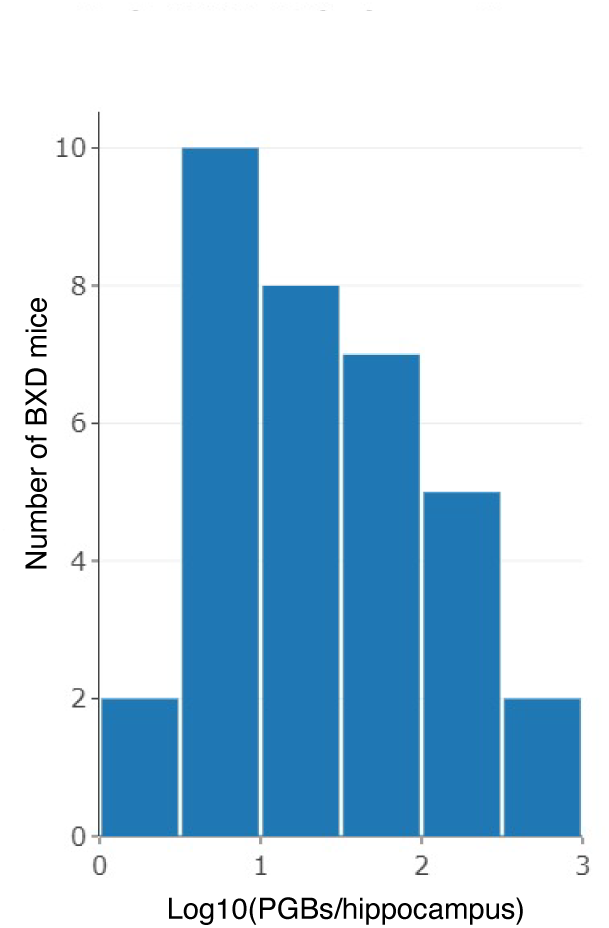
Log transformed PGB count distribution. Polyglucosan accumulation in the hippocampus shows a normal log-transformed distribution across BXD strains.

**Figure S2.**
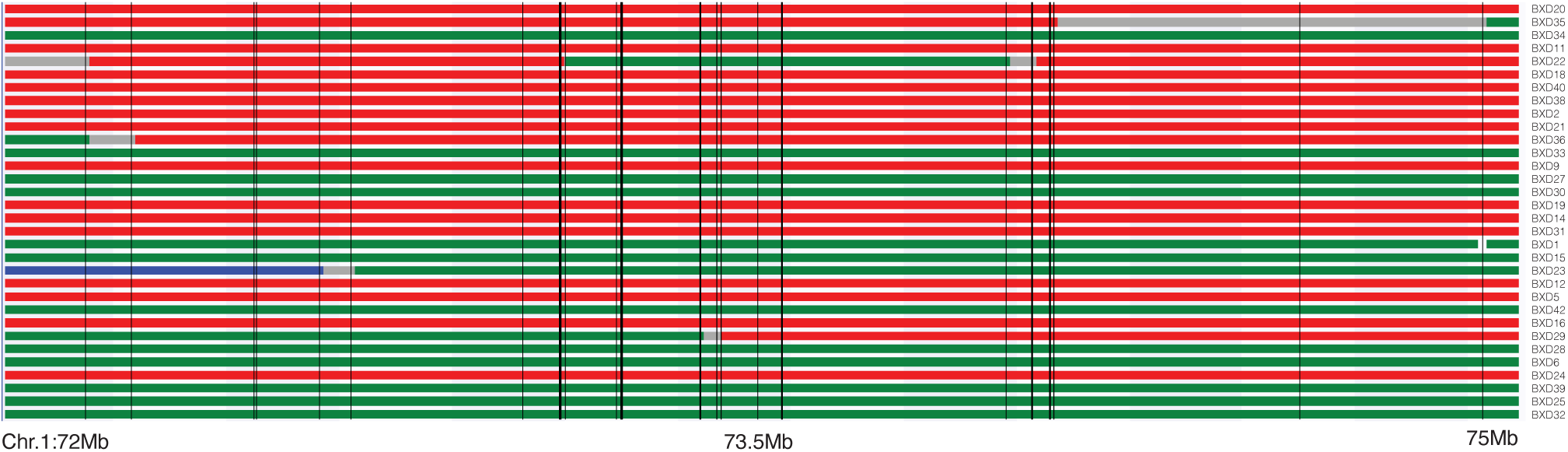
Heterogenous allele inheritance from the DBA/2J and C57BL/6J strains at the *Pgb1* locus across the BXD family. Inheritance from either the *B* (red) or *D* (green) parental allele within the *Pgb1* locus. Blue indicates a heterozygous carrier. Chr. = Chromosome, Mb = megabases.

**Figure S3.**
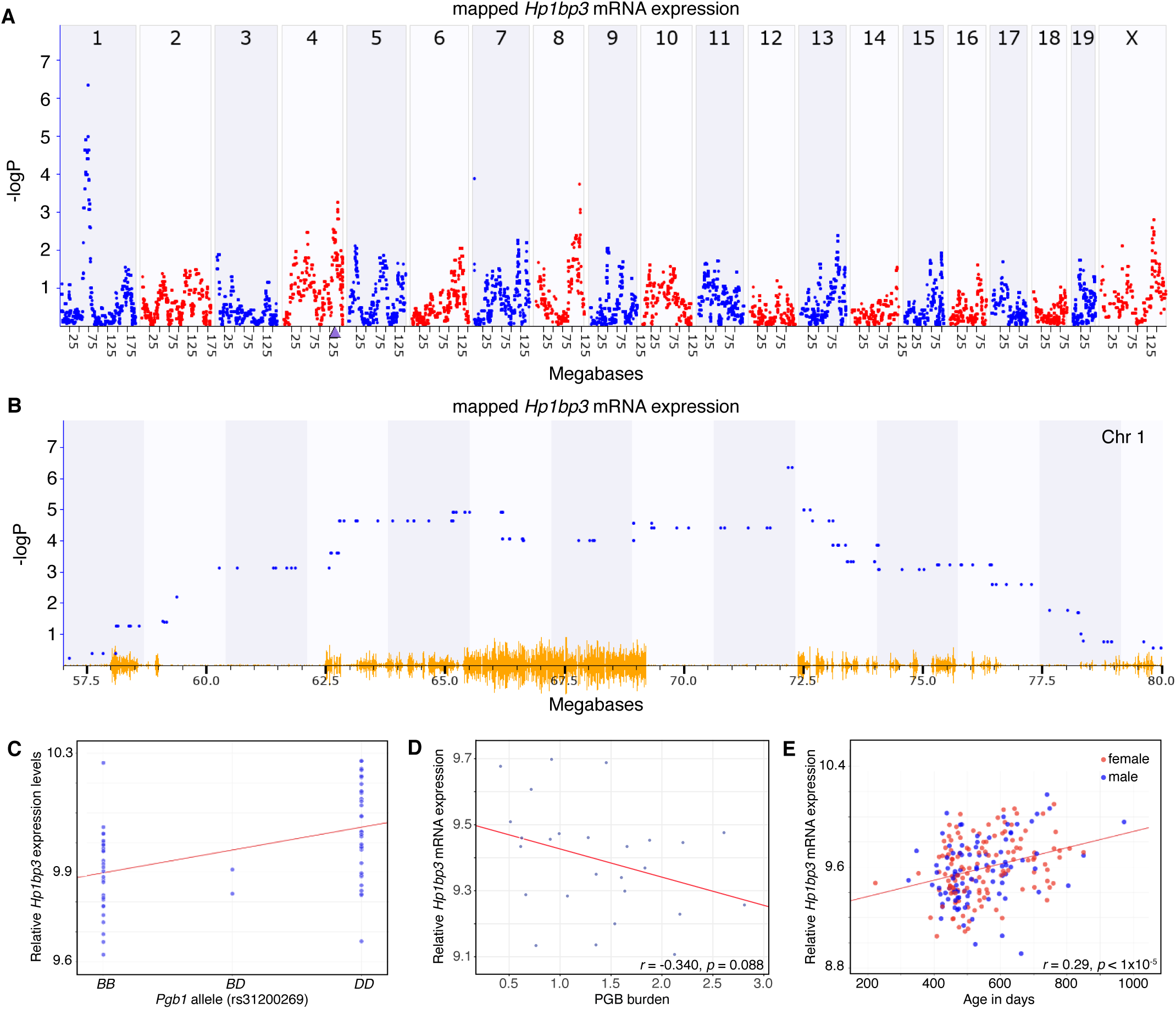
*Hp1bp3* mRNA expression is regulated by the *Pgb1* locus. (**A**) The *Hp1bp3* transcript maps as a *trans-*eQTL to *Pgb1* on Chr 1 at ~72.2 Mb. Note that there is a *cis*-eQTL at the location of the cognate *Hp1bp3* gene on Chr 4 at 138.2 Mb (purple triangle on x-axis). (**B**) Enlarged view of the *trans*-eQTL peak location. The highest −logP value is ~300 Kb downstream of *Smarcal1* at 72.54 Mb. Yellow indicates allele variations annotated at GeneNetwork.org. (**C**) Scatterplot of the expression of *Hp1bp3* as a function of *B* and *D* alleles highlighting the protective effect of the *D* allele. (**D**) Scatterplot indicating the direct correlation between *Hp1bp3* mRNA expression and PGB density. (**E**) Scatterplot of the expression of *Hp1bp3* as a function of age.

**Figure S4.**
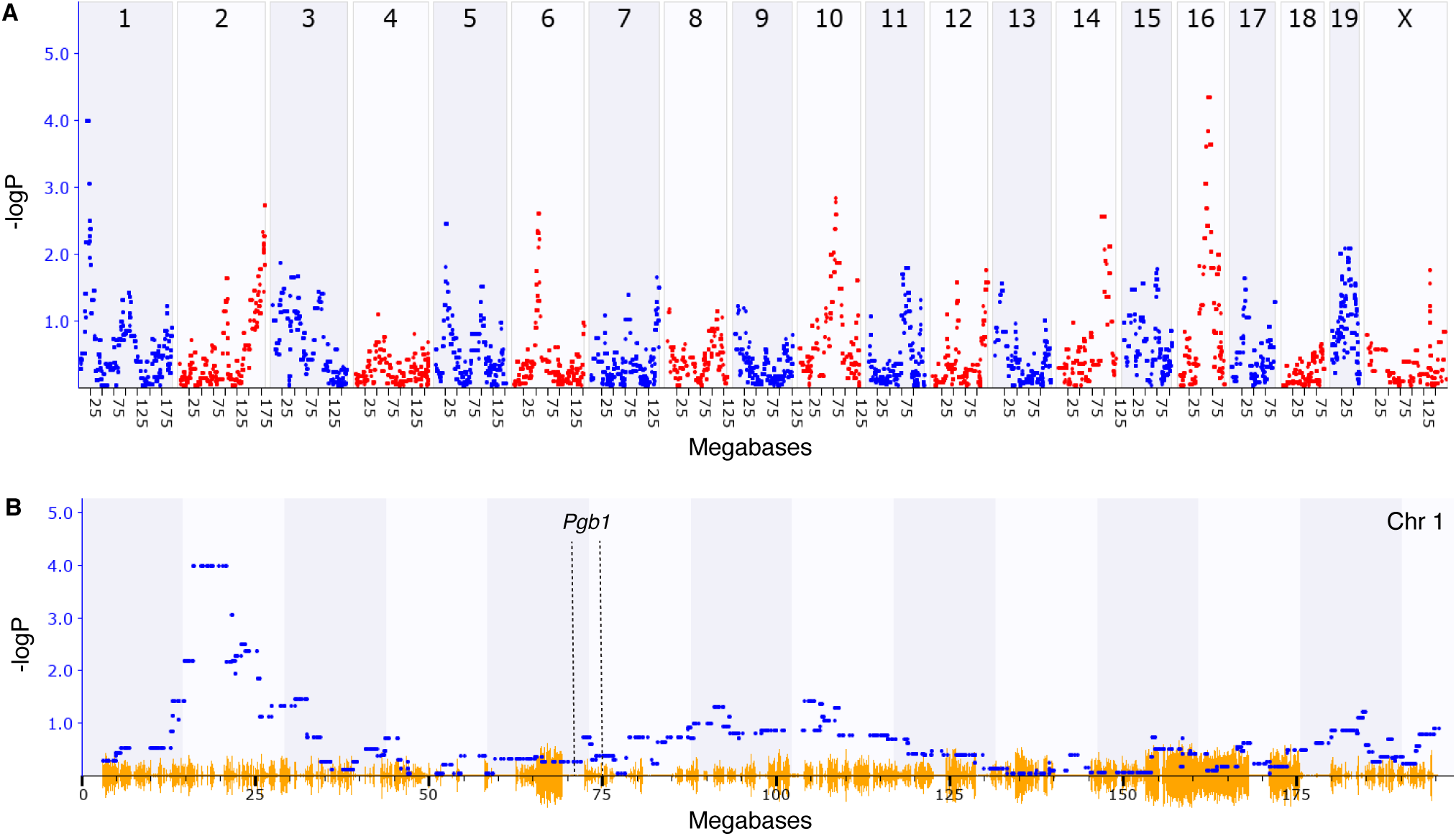
Conditioned fear memory performance does not map to the *Pgb1* locus. (**A**) Mapping of the conditioned fear memory performance using GEMMA shows two major correlating loci (-logP ≥ 4) on chromosome 1 and chromosome 16. (**B**) No correlation was found between conditioned fear memory performance and the *Pgb1* locus on chromosome 1.

**Figure S5.**
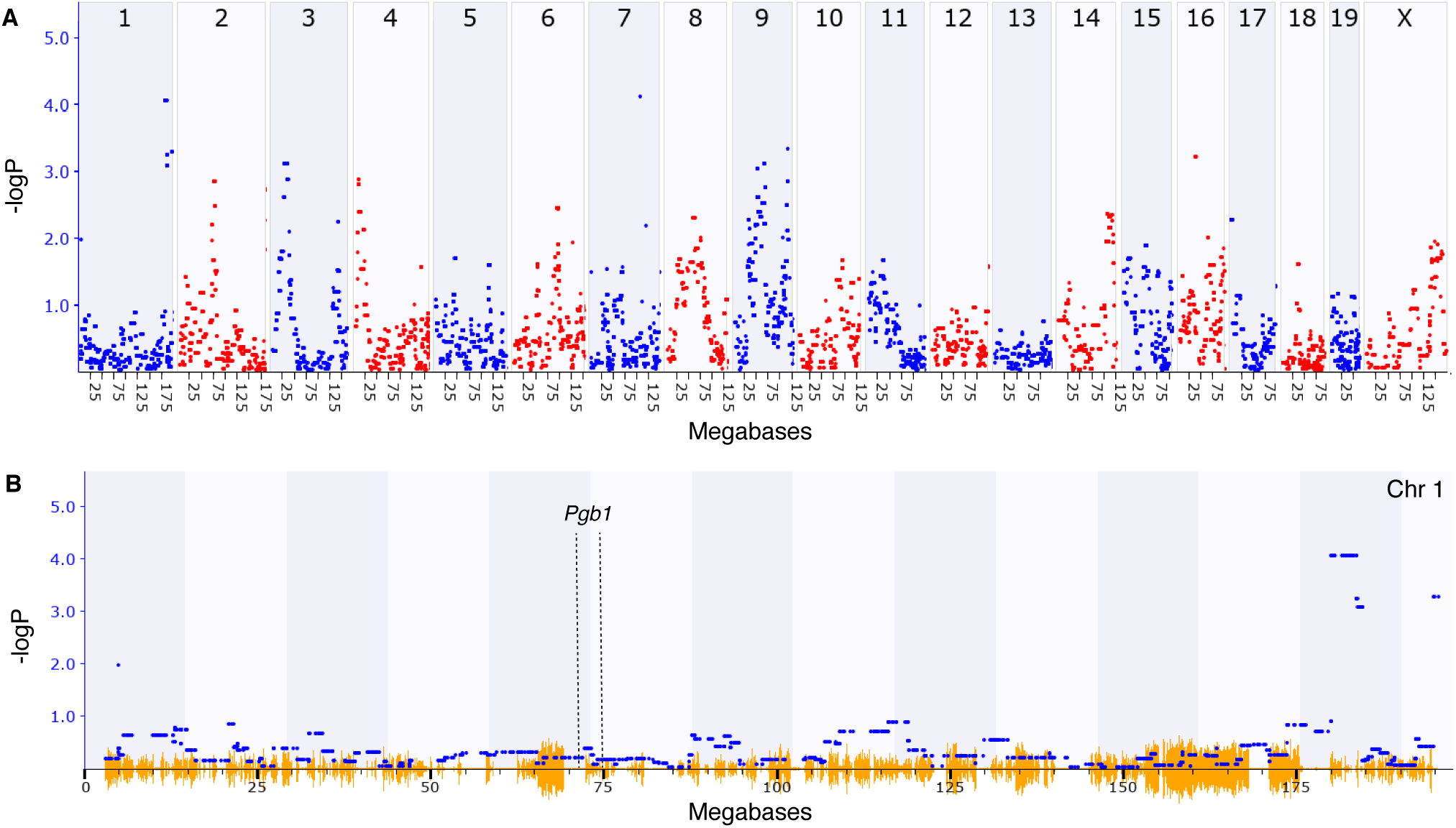
Y-maze performance does not map to the *Pgb1* locus. (**A**) Mapping of the Y-Maze performance using GEMMA shows two major correlating loci (-logP ≥ 4) on chromosome 1 and chromosome 7. (**B**) No correlation was found between Y-maze performance and the *Pgb1* locus on chromosome 1.

